# Liquid-Liquid Phase Separation Protects Amyloidogenic and Aggregation-Prone Peptides in Heterologous Expression Systems

**DOI:** 10.1101/2021.05.14.443401

**Authors:** Bartosz Gabryelczyk, Margaret Philips, Kimberly Low, Anandalakshmi Venkatraman, Bhuvaneswari Kannaian, Reema Alag, Markus Linder, Konstantin Pervushin, Ali Miserez

## Abstract

Studying pathogenic effects of amyloids requires homogeneous amyloidogenic peptide samples. Recombinant production of these peptides is challenging due to their susceptibility to aggregation and chemical modifications. Thus, chemical synthesis is primarily used to produce amyloidogenic peptides suitable for high resolution structural studies. Here, we exploited the shielded environment of protein condensates formed via liquid-liquid phase separation (LLPS) as a protective mechanism against premature aggregation. We designed a fusion protein tag undergoing LLPS in *E. coli* and linked it to highly amyloidogenic peptides, including Aβ amyloid. We find that the fusion proteins form membraneless organelles during overexpression and remain soluble. We also developed a facile purification method of functional Aβ peptides free of chromatography steps. The strategy exploiting LLPS can be applied to other amyloidogenic, hydrophobic, and repetitive peptides that are otherwise difficult to produce.

---

Cost-effective and high-yield production of amyloidogenic peptides, associated with several neuropathologies,^[1]^ remains an important challenge for their detailed structural and *in vitro* investigations. For example, NMR-based spectroscopic studies require isotopically-labelled peptides to obtain their high-resolution three-dimensional structures.^[2,3]^ There are currently two main methods used for amyloidogenic peptide production, namely solid-phase peptide synthesis (SPPS)^[4]^ and recombinant expression.^[5]^ SPPS is usually a straightforward way to obtain many peptides. However, synthesis of aggregation-prone and amyloidogenic sequences is often challenging due to their high hydrophobicity and self-assembling tendencies.^[6]^ Therefore, it is usually difficult and very costly to produce sufficient quantities of these peptides, especially isotopically labelled.

In contrast, recombinant expression of amyloidogenic peptides in *Escherichia coli* (*E. coli*) is less expensive, more suitable for large scale production, and can be combined with isotope labelling.^[5]^ However, high yield production of peptides is affected by their inherent propensity to aggregate and associated toxicity.^[2]^ These challenges can potentially be solved by the formation of insoluble inclusion bodies in the host cells.^[7]^ Nevertheless, extraction of the peptides from the inclusion bodies is often time-consuming, require use of harsh denaturing agents, and may lead to only partial refolding of purified peptides, thus affect their biologically-active conformations.^[8]^ Hence, peptides recovered from inclusion bodies may not be ideal for highly sensitive NMR studies.

Solubility tags may prevent aggregation of recombinant proteins during expression, allowing the use of purification methods under non-denaturing conditions.^[9]^ However, tags must be cleaved off from the fusion partner after protein expression, requiring an additional purification step. The cleavage is usually achieved by proteolytic enzymes or chemically induced and requires specific reaction conditions that may sometimes lead to chemical modifications of the target peptide.

To overcome these challenges, it is desirable to produce recombinant amyloidogenic peptides that do not prematurely aggregate in the host cell, while at the same time preventing their sequestration in inclusion bodies. In this study, we hypothesized that this could be achieved by fusing amyloid peptides to a peptide domain that rapidly forms membraneless organelles within the host cells *via* liquid-liquid phase separation (LLPS). Membraneless organelles are highly concentrated microdroplets that phase-separate from a homogenous solution of biomolecules.^[10,11]^ Unlike inclusion bodies, concentrated macromolecules within these organelles are not aggregated but maintain a high mobility,^[12]^ resulting in liquid-like properties. Initially discovered in eukaryotic cells,^[10]^ membraneless organelles have recently been achieved synthetically in *E. coli*.^[13]^ We reasoned that by forming liquid condensates, the expressed fusion proteins would bury the hydrophobic amyloidogenic domains and prevent early aggregation. To achieve this, we designed a fusion tag derived from histidine-rich beak protein 1 (HBP-1), which we recently found to exhibit single-component LLPS.^[14,15]^

As model peptides, we selected sequences that are notoriously difficult to produce with both SPPS and recombinant expression methods, *i.e*.: *(i)* two peptides from the highly amyloidogenic regions of the transforming growth factor β-induced (TGFBI) protein associated with corneal dystrophies,^[16,17]^ which were previously shown to be highly insoluble in an *E. coli*-based expression;^[18]^ and *(ii)* an engineered variant of Aβ 17-40 peptide (Aβ-P3*) associated with Alzheimer’s disease.^[19]^ Aβ-P3 exhibits faster kinetics of fibril formation than other full-length Aβ peptides,^[20]^ resulting in increased cytotoxicity in cultured neuronal cells.^[21]^ Because it lacks the N-terminus region of other Aβ peptides that helps maintain partial solubility,^[22]^ Aβ-P3 peptide is very unstable, resulting in accelerated aggregation and self-assembly of fibrils almost immediately after production.^[20]^ Using fluorescence imaging, we find that following overexpression the fusion constructs form membraneless organelles in *E. coli*, subsequently facilitating their purification while at the same time preventing premature aggregation. Once the LLPS-forming tag is cleaved off, the amyloidogenic peptides spontaneously self-assemble into amyloid fibrils amenable to structural studies as shown from TEM observations. Additional hydrophobic and repetitive sequences that have been challenging to express in soluble form were also produced to broaden the applicability of the tag.

In a recent study we showed that truncated HBP-1 protein variants can undergo LLPS in a similar way as the full-length protein. We also found that a specific sequence motif from the repetitive modular sequence of the C-terminal region of the protein drives LLPS.^[15]^ Since the N-terminal domain of the protein does not contribute to LLPS, we speculated that its role might be to facilitate expression of the repetitive and hydrophobic –and thus aggregation-prone– C-terminal sequence. Using this knowledge, we designed a HBP-1-derived sequence called BEAK-tag composed of the (non-repetitive) N-terminal region of HBP-1 and part of the repetitive modular domain (responsible for initiating LLPS) (Figure S1). We then tested whether it could be used to facilitate expression of amyloidogenic peptides and other repetitive and hydrophobic sequences (Figure S2, Figure 1).

**Figure 1.**
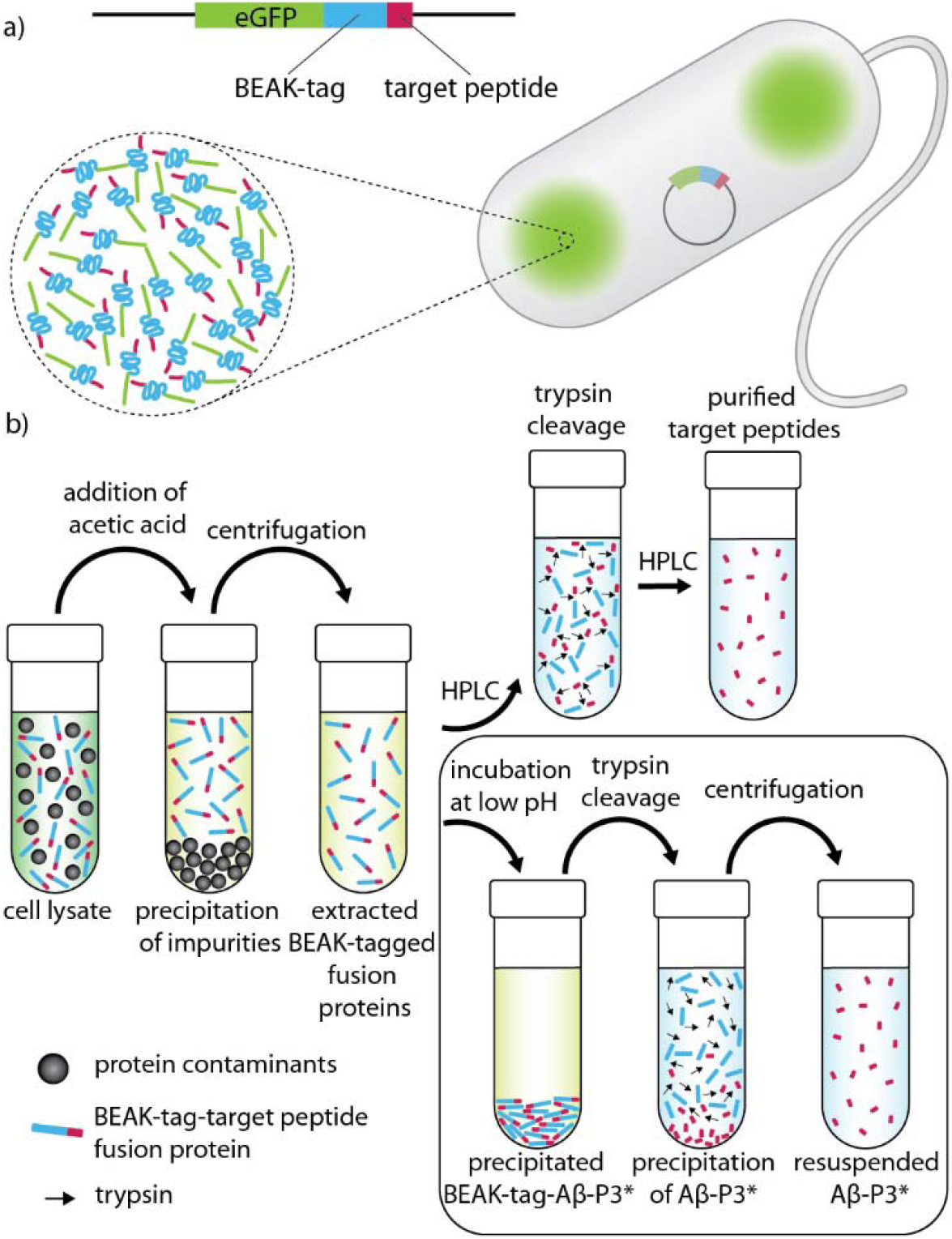
Schematic expression and purification process of BEAK-tagged amyloidogenic peptides. **a)** Expression of the fusion proteins leads to the formation intracellular condensates. Proteins within the condensates are highly concentrated but remain fluidic due to BEAK-tag properties, thus preventing premature aggregation of the amyloidogenic peptide. **b)** Purification protocol based on low pH extraction form the soluble fraction. The Aβ-P3*peptide can be obtained without additional chromatography purification (as described in the lower-right box).

Fluorescent microscopy was used to monitor protein expression inside bacterial cells. Approximately 1 – 2 hours after inducing protein expression, we observed the appearance of condensed structures in the polar regions of the bacterial cells for all constructs under study. Very similar structures have recently been observed in *E. coli* for silk- ^[13]^ and resilin- ^[23]^ like proteins and were described as protein condensates typically observed in membraneless organelles.

In order to investigate whether the observed structures in our experiments exhibited liquid-like properties that are typical for membraneless organelles, we mixed the bacterial cells with a lysis buffer containing lysozyme, a detergent that disrupts the cell membrane. Incubation of bacterial cells in the lysis buffer disrupted the cells’ structures and caused diffusion of cytoplasm into the surrounding buffer. At this point, we also observed that the fluorescence signal from the protein condensates diffused, indicating that they were also disassembled (Figure 2b and Figure S3). Thus, the condensates exhibited liquid-like properties and their constitutive proteins were soluble and not randomly aggregated as in the case of inclusion bodies. These results confirmed our hypothesis that the BEAK-tag fusion proteins, upon overexpression in bacterial cells, assembled in protein condensates resembling membraneless organelles with their proteins having significant molecular mobility (Figure 1a).

**Figure 2.**
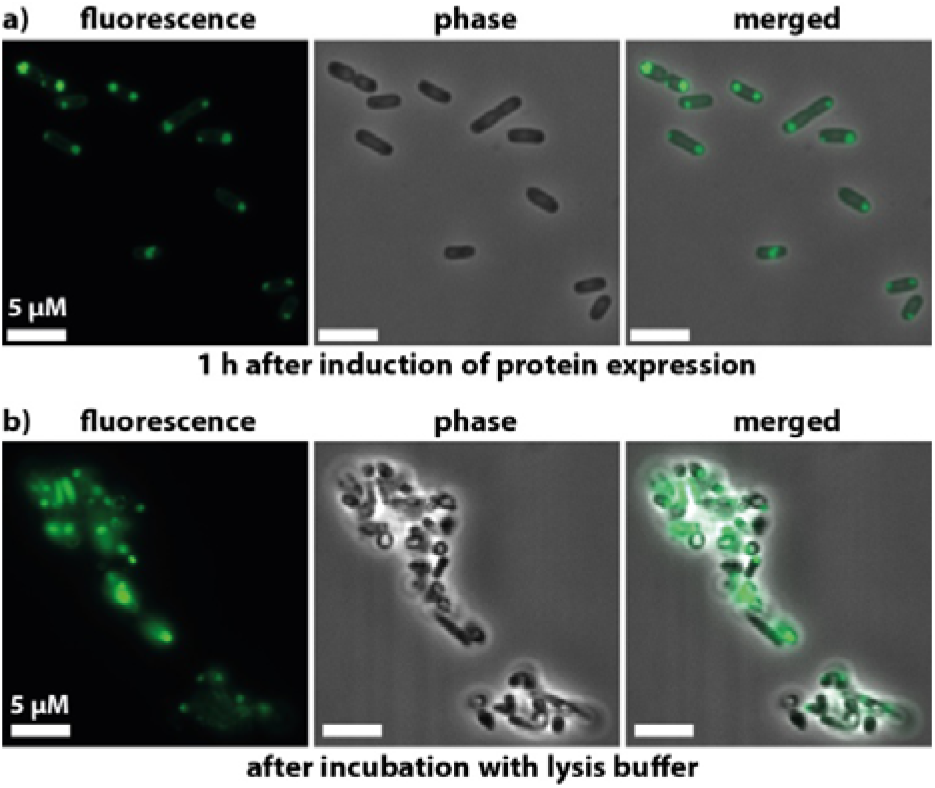
Fluorescence microscopy and phase contrast images of bacterial cells expressing eGFP-BEAK-tag-TGFBPIp1 fusion protein. a) Fluorescent condensates localized near the polar regions of the cells indicate the formation of membraneless organelles. b) Diffusion of the fluorescent signal indicating liquid-like properties of the protein condensates.

Next, we expressed the BEAK-tag fusion proteins (without eGFP) and analyzed their solubility after cell lysis. All fusion proteins were found in the soluble fraction (Figure S4) and were then extracted using a low pH by addition of concentrated acetic acid (Figure 1b). The pH change of the soluble fraction (form pH 8.0 of the lysis buffer to approximately pH 3.3 after the addition of acetic acid) caused rapid precipitation of most bacterial proteins and other contaminants released from bacterial cells during lysis. However, it did not affect the BEAK-tagged proteins, which remained fully soluble (Figure S4). The high solubility of the BEAK-tag at low pH was especially beneficial to isolate the very hydrophobic Aβ-P3* peptide. Since the aggregation of the Aβ peptides is pH-dependent and the fastest aggregation rates were observed at pH 4.5 to 9,^[24]^ using low pH in combination with the BEAK-tag allowed to maintain it in solution in the initial step of the purification process. This also enabled us to use centrifugation to remove precipitated impurities and obtain a solution of relatively pure BEAK-tag-Aβ-P3* fusion protein (Figure S4c). Eventually, incubation of the protein in a low pH solution led to its precipitation while the remaining impurities from the cell lysate (soluble in acidic pH) remained in solution. The aggregated fusion protein was then collected by centrifugation and the protein pellet re-dissolved in NaOH solution. In that way, the purity of the BEAK-tag-Aβ-P3* fusion protein was further increased and did not require an additional purification step. On the other hand, the extracted BEAK-tagged TGFBIp and repetitive peptides remained fully soluble at low pH and thus required a final purification step with reverse phase HPLC (Figure S4a,b).

NMR was used to further verify the solubility and integrity of the BEAK-tag-TGFBIp fusion proteins after purification. Figure S5 presents an overlay of the ^1^H-^15^N-HSQC NMR spectrum of the BEAK-tag (without fused peptide) and the BEAK-tag-TGFBIp fusion proteins. The spectra showed the presence of well-resolved cross-peaks that could be readily assigned to the TGFBIp peptides, thus confirming that the fusion protein was soluble.

Next, the purified fusion proteins were subjected to trypsin cleavage in order to obtain the target peptides. Since the solubility of the BEAK-tag is also high at basic pH^[15]^, the cleavage reaction could be conducted under optimal pH for trypsin activity. This allowed to obtain pure Aβ-P3* peptide without any further purification step. During the cleavage reaction, the concentration of the free peptide Aβ-P3* gradually increased leading to its rapid aggregation, while the released BEAK-tag remained in solution. As a result, the precipitated Aβ-P3* could be collected by centrifugation and then dissolved in NaOH solution. On the other hand, TGFBIp and the repetitive peptides did not aggregate after cleavage and required purification with HPLC. The MW of purified amyloidogenic peptides was verified by MALDI-TOF mass spectroscopy (Figure S6).

Finally, we tested whether the possible limitations of our expression and purification methods could be circumvented using alternative procedures. For example, the use of trypsin for the cleavage reaction requires that a target peptide does not harbor any Lys or Arg residues, while low pH conditions may affect sensitive target peptides. Thus, we prepared a control construct (His-BEAK-tag-TEV-TGFBp2), in which an additional His-tag was fused to the N-terminus of the BEAK-tag and the trypsin recognition site was substituted with Tobacco Etch Virus (TEV), respectively. We then purified the construct using affinity chromatography and successfully cleaved the target peptide form the BEAK-tag with TEV protease (Figure S7).

Purified amyloidogenic peptides were tested to assess whether they were able to stably and spontaneously form amyloid fibers. Investigations with transmission electron microscopy (TEM) (Figure 3) clearly showed that all tested peptides formed homogeneous amyloid fibrils with an apparent morphology corresponding to that observed in their SSPS counterparts. TGFBIp1 showed long fibrils up to > 200 nm in length within one week of growth (Figure 3a). TGFBIp2 (Figure 3b) formed shorter fibrils compared to TGFBIp1. Structural polymorphisms cannot be assessed at the resolution provided by TEM and will require further cryoEM investigations. Fibrils formed by Aβ-P3*peptides (Figure 3c) exhibited a diameter of approximately 20 nm and twists in their ribbon-like features, which is consistent with previously observed TEM images of various types of Aβ fibrils.^[20,25]^

**Figure 3.**
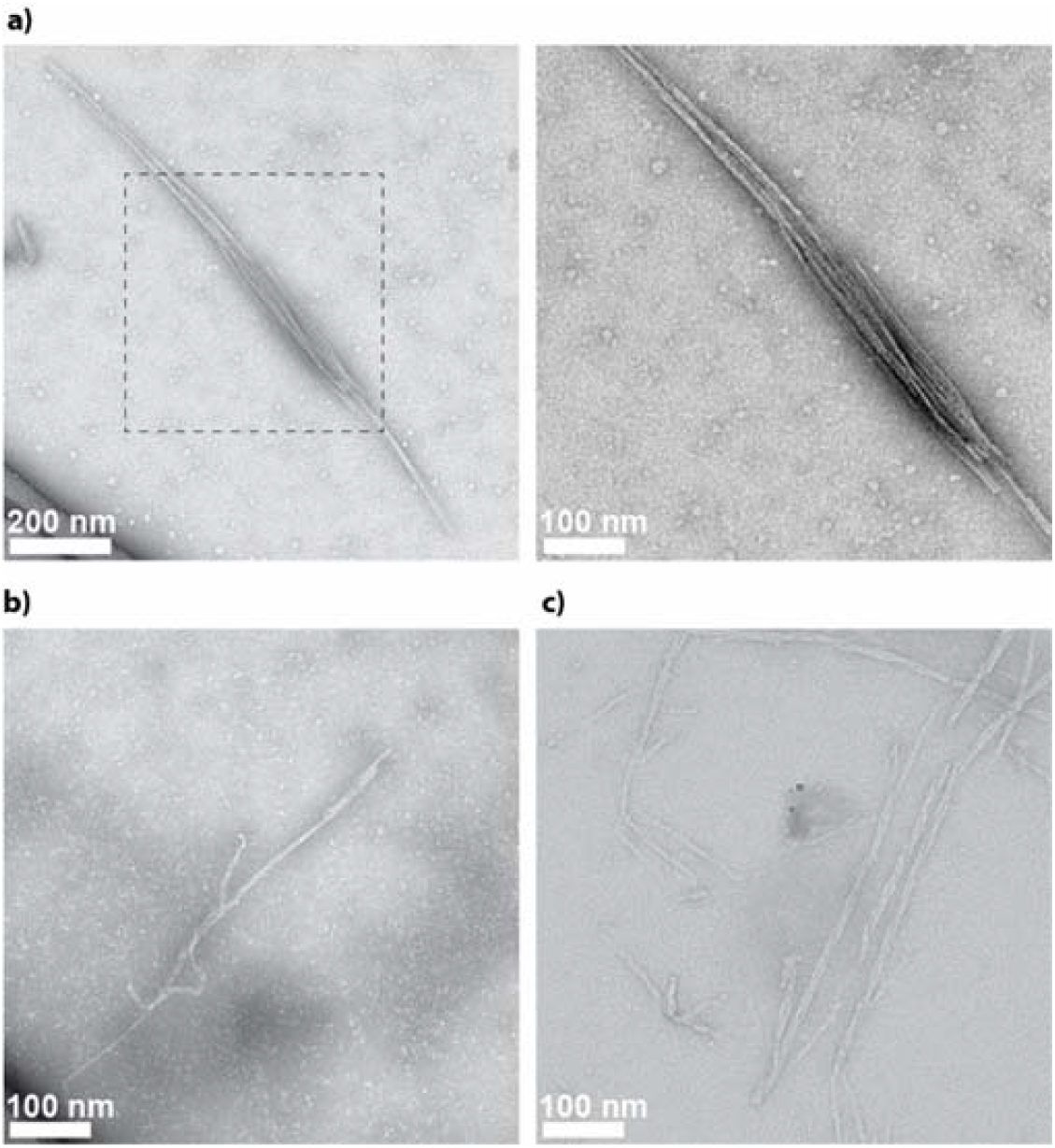
TEM images of the amyloid fibrils formed by amyloidogenic peptides. a) TGFBIp1 at low (left) and high (right) magnification. b) TGFBIp2. c) Aβ-P3*.

Overall, our data indicate that LLPS can be exploited to efficiently produce amyloidogenic and hydrophobic peptides that are difficult to obtain *via* SSPS and recombinant expression. Following overexpression in *E. coli*, the BEAK-tag fusion proteins formed protein condensates resembling membraneless organelles found in eukaryotic cells.^[10,11]^ The assembly of similar protein condensates in *E. coli* has very recently been reported in other proteins with LLPS properties, including silk-^[13]^ and resilin-like^[23]^ proteins. Our results are in line with these studies and show that the protein condensates exhibit liquid-like properties. The proteins in the condensates are highly concentrated but still fluidic and not aggregated as in the case of insoluble inclusion bodies. These results indicate that the hydrophobic sequences within the protein condensates might undergo some molecular arrangement that prevent them from aggregating.

BEAK-tag not only prevents protein aggregation *in vivo* but also serves as a solubility tag, thereby facilitating purification. Hence, any undesirable chemical modification or incomplete refolding of the peptides (as in the case of insoluble expression into inclusion bodies) is minimised. In addition, trypsin cleavage enables very efficient enzymatic digestion of target peptides from the fusion tag. This simplified procedure is notably related to the absence of any basic residues in the BEAK-tag. This potential limitation can be circumvented by engineering the cleavage site. Indeed, our data show that fully functional peptides can also be obtained when TEV protease was used instead of trypsin. Our approach can therefore be used to obtain pure, very hydrophobic sequences such as Aβ-P3* without the need for additional chromatography purification steps, offering significant advantages compared to other recombinant expression methods of Aβ peptides that often involve long and complex procedures.^[2]^

In summary, this proof-of-concept study shows that BEAK-tag can be used to efficiently produce amyloidogenic peptides by exploiting LLPS within *E. coli*, which shields the target peptides from premature aggregation. Our expression and purification methods are inexpensive, fast, and lead to high yield of functional amyloidogenic peptides. The BEAK-tag can also be used to facilitate expression of a range of aggregation-prone sequences, including other fragments of amyloid beta and hydrophobic modular repetitive peptides that are otherwise challenging to produce using current recombinant expression and SPPS protocols.

## Acknowledgements

This research was funded by the Singapore Ministry of Education (MOE) through an Academic Research Fund (AcRF) Tier 3 grant (grant # MOE 2019-T3-1-012) and the Academy of Finland project 315140.

## References

[1] F. Chiti, C. Dobson, Annu. Rev. Biochem. 2017, 86, 27–68.

[2] L. Jia, W. Zhao, W. Wei, X. Guo, W. Wang, Y. Wang, J. Sang, F. Lu, F. Liu, Crit. Rev. Biotechnol. 2020, 40, 475–489.

[3] A. Loquet, N. El Mammeri, J. Stanek, M. Berbon, B. Bardiaux, G. Pintacuda, B. Habenstein, Methods 2018, 138-139, 26–38.

[4] L. Raibaut, O. El Mahdi, O. Melnyk, Top. Curr. Chem. 2015, DOI 10.1007/128_2014_609.

[5] S. Wegmuller, S. Schmid, Curr. Org. Chem. 2014, 18, 1005–1019.

[6] M. Paradís-Bas, J. Tulla-Puche, F. Albericio, Chem. Soc. Rev. 2016, 45, 631–654.

[7] P. M. Hwang, J. S. Pan, B. D. Sykes, FEBS Lett. 2014, 588, 247–252.

[8] H. Yamaguchi, M. Miyazaki, Biomolecules 2014, 4, 235–251.

[9] G. L. Rosano, E. A. Ceccarelli, Front. Microbiol. 2014, 5, 172.

[10] S. F. Banani, H. O. Lee, A. A. Hyman, M. K. Rosen, Nat. Rev. Mol. Cell Biol. 2017, 18, 285–298.

[11] I. Peran, T. Mittag, Curr. Opin. Struct. Biol. 2020, 60, 17–26.

[12] D. Bracha, M. T. Walls, C. P. Brangwynne, Nat. Biotechnol. 2019, 37, 1435–1445.

[13] S.-P. Wei, Z.-G. Qian, C.-F. Hu, F. Pan, M.-T. Chen, S. Y. Lee, X.-X. Xia, Nat. Chem. Biol. 2020, 16, 1143–1148.

[14] H. Cai, B. Gabryelczyk, M. S. S. Manimekalai, G. Grüber, S. Salentinig, A. Miserez, Soft Matter 2017, 13, 7740–7752.

[15] B. Gabryelczyk, H. Cai, X. Shi, Y. Sun, P. J. M. Swinkels, S. Salentinig, K. Pervushin, A. Miserez, Nat. Commun. 2019, 10, 5465.

[16] A. Venkatraman, B. Dutta, E. Murugan, H. Piliang, R. Lakshminaryanan, A. C. Sook Yee, K. V. Pervushin, S. K. Sze, J. S. Mehta, J. Proteome Res. 2017, 16, 2899–2913.

[17] V. Anandalakshmi, E. Murugan, E. G. T. Leng, L. W. Ting, S. S. Chaurasia, T. Yamazaki, T. Nagashima, B. L. George, G. S. L. Peh, K. Pervushin, R. Lakshminarayanan, J. S. Mehta, Biochem. J. 2017, 474, 1705–1725.

[18] M. Elavazhagan, R. Lakshminarayanan, L. Zhou, L. W. Ting, L. Tong, R. W. Beuerman, S. S. Chaurasia, J. S. Mehta, Protein Expr. Purif. 2012, 84, 108–115.

[19] W. Wei, D. D. Norton, X. Wang, J. W. Kusiak, Brain 2002, 125, 2036–2043.

[20] A. J. Kuhn, B. S. Abrams, S. Knowlton, J. A. Raskatov, ACS Chem. Neurosci. 2020, 11, 1539–1544.

[21] H. S. Nhan, K. Chiang, E. H. Koo, Acta Neuropathol. 2015, 129, 1–19.

[22] C. Morris, S. Cupples, T. W. Kent, E. A. Elbassal, E. P. Wojcikiewicz, P. Yi, D. Du, Chem. - A Eur. J. 2018, 24, 9494–9498.

[23] M. Dzuricky, B. A. Rogers, A. Shahid, P. S. Cremer, A. Chilkoti, Nat. Chem. 2020, 12, 814–825.

[24] S. Kobayashi, Y. Tanaka, M. Kiyono, M. Chino, T. Chikuma, K. Hoshi, H. Ikeshima, J. Mol. Struct. 2015, 1094, 109–117.

[25] M. Schmidt, C. Sachse, W. Richter, C. Xu, M. Fändrich, N. Grigorieff, Proc. Natl. Acad. Sci. USA. 2009, 106, 19813–19818.

