## Supplementary Information for "Liquid-Liquid Phase Separation Protects Amyloidogenic and Aggregation-Prone Peptides in Heterologous Expression Systems"

**Table of contents**

EXPERIMENTAL DETAILS 3

Design of BEAK-tag fusion proteins 3

Protein expression 3

Enzymatic cell lysis 4

Fluorescence microscopy 4

Extraction of BEAK-tagged fusion proteins 4

Reverse-phase HPLC 4

Trypsin cleavage 5

Matrix assisted laser desorption/ionization-time of flight (MALDI-TOF) mass spectroscopy 5

Sodium dodecyl sulphate polyacrylamide gel electrophoresis (SDS-PAGE) 5

NMR 5

TEM sample preparation and imaging 6

Thioflavin T (ThT) assay 6

FIGURES 7

Figure S1. BEAK-tag schematic illustration of the construct. 7

Figure S2. Protein and peptide sequences used in the study. 8

Figure S3. Fluorescence microscopy and phase contrast images of bacterial cells expressing. 9

Figure S4. BEAK-tag facilitates recombinant production of biologically functional aggregation-prone peptides. 10

Figure S5. Overlay of the 1H-15N-HSQC NMR spectra of the BEAK-tag (without fused peptide) 11

Figure S6. MALDI-TOF spectra of purified amyloidogenic peptides 12

Figure S7. Qualitative and functional analysis of the TGFBIp2 peptide obtain after TEV cleavage of the BEAK-tag-TEV-TGFBIp2 control protein. 13

### EXPERIMENTAL DETAILS

#### Design of BEAK-tag fusion proteins

BEAK-tag fusion proteins constructs were created by combining BEAK-tag *i.e.* first 112 amino acids of HBP-1 protein (UniProtKB - A0A0G2UMW8, with methionine at the position 1 instead of the signal peptide) with lysine residue, followed by a target peptide sequence (Figure S1). Amino acid sequences of the constructs are presented in Figure S2. Synthetic genes encoding BEAK-tag fusion proteins were obtained from ATUM (USA). The constructs were sub-cloned into pJ414 expression vector (containing inducible T7 promoter) and then transformed into chemically competent *E. coli* BL21 (DE3) cells.

eGFP-BEAK-tag-peptide constructs with the enhanced green fluorescent protein (eGFP, used for fluorescence microscopy studies, Figure 2 and S3) were assembled using the golden gate cloning[1] and subcloned into pET-28a expression vector. The plasmids were then transformed into chemically competent BL21A1 *E. coli* strain (Thermo Fisher).

BEAK-tag-TEV-TGFBIp2 construct (used for control experiments, Figure S7) containing Tobacco Etch Virus (TEV) protease cleavage inserted between BEAK-tag and TGFBIp2 peptide was obtained from Bio Basic (Singapore). The synthetic gene was sub-cloned into pET-15b expression vector downstream of N-terminal His tag and transformed into *E. coli* BL21 (DE3) cells.

#### Protein expression

eGFP-BEAK-tag-(target peptide) fusion proteins for florescence microscopy experiments. The *E. coli* cells containing expression plasmids with eGFP-BEAK-tag-(target peptide) fusion protein genes were cultivated in LB medium (2 mL) supplemented with 0.2 % glucose and 50 µg/mL kanamycin. Bacterial cultures were inoculated with a single colony from LB-agar plates and incubated at 37° C, 220 RPM, overnight (approximately 16 – 18 hours). Next, the overnight cultures were diluted 1/50 and incubated until $OD_{600}$ reached 0.5. Protein expression was induced with 1 mM IPTG and 0.2% L-arabinose (final concentration).

BEAK-tag-(target peptide) fusion proteins for solubility studies, NMR, TEM. The *E. coli* cells containing expression plasmids with BEAK-tag-(target peptide) fusion protein genes were cultivated in M9 medium supplemented with 0.4 % glucose, 1 mg/L biotin, 1 mg/L thiamine, and 100 µg/mL ampicillin. The precultures were incubated overnight at 37° C, 220 RPM and then diluted 1/100 and used to inoculate main cultures. Protein expression was induced with 0.5 mM IPTG (final concentration) at $OD_{600}$ = 0.5 – 0.6 and carried out overnight at 37 °C, 220 RPM. Cells were then harvested by centrifugation (8000 RPM, 10 min, 4 °C) and lysed immediately or stored at -20 °C.

BEAK-tag-TEV-TGFBIp2 for control experiments. The *E. coli* cells containing expression plasmid with BEAK-tag-TEV-TGFBIp2 were grown in LB media containing 100 µg/mL ampicillin. Protein expression was induced using 0.5 mM IPTG (final concentration) when $OD_{600}$ reached between 0.5 – 0.6 and was carried out for 4 hours at 37° C, 220 RPM. Cells were collected by centrifugation and resuspended in a lysis buffer containing 20 mM Tris pH 8.0, 1mM EDTA, 300 mM NaCl and protease inhibitors (Calbiochem). Cell lysis was performed using sonication. Lysate was centrifuged at 18000 RPM for 20 min at 4 °C. Supernatant containing the fusion protein was then passed through $Ni^{2+}$-NTA resin. The protein was eluted from the resin with the buffer containing 20 mM Tris pH 8.0, 300 mM NaCl and 500 mM imidazole. Buffer exchange to the final TEV cleavage buffer: 20 mM Tris pH 8, 200 mM NaCl and 2 mM TCEP was performed on PD 10. TGFBIp2 peptide was cleaved from the BEAK-tag using TEV protease with 1:100 mass ratio (TEV protease/fusion protein) at 4 °C overnight. The reaction was then passed through $Ni^{2+}$-NTA resin again. Cleaved peptide was collected in flow through while the BEAK-tag remained bound to the resin. Additional, reverse phase-HPLC (details described below) was performed to further increase the purity of the peptide.

**Enzymatic cell lysis**

Bacterial cells expressing eGFP-BEAK-tag-(target peptide) fusion proteins were harvested by centrifugation (5000 RPM, 3 min, 21 °C) 1 – 2 hours after induction of protein expression. Cells were then resuspended in a lysis buffer composed of 10 mM Tris-HCl pH 8.0, 1 mM EDTA, 1% Triton X-100, 2 mg/ml lysozyme (L6876 Sigma-Aldrich) and incubated 30 min before fluorescence microscopy imaging.

**Fluorescence microscopy**

Fluorescence and phase contrast and images were acquired using an Axio Observer Z1 microscope (Carl Zeiss, Jena, Germany) at 100x magnification. GFP signal was obtained using excitation light at 480 nm, while collecting the emitted light of 515 – 535 nm. Samples were prepared onto a standard microscope slides coated with Poly-D-Lysine (A3890401 Thermo Fisher).

**Extraction of BEAK-tagged fusion proteins**

Cells pellets containing overexpressed fusion proteins were resuspended in a lysis buffer composed of 20 mM Tris pH 8.0, 1 mM EDTA, and cOmplete™ Protease Inhibitor Cocktail (Roche). Cell lysis was performed with a microfluidizer (18000 psi, 5 passes, 4 °C). Lysate was centrifuged (15000 RPM, 15 min, 4 °C) immediately after cell lysis. Next, soluble fraction (supernatant) was mixed with ice cold 100 % acetic acid in a volume ratio of 1:20 (acetic acid/cell lysate), homogenized by gentle vortexing, and centrifuged (15000 RPM, 10 min, 4 °C). The pellet (containing precipitated impurities) was discarded while supernatant (containing fusion protein) of the BEAK-tag-TGFBIp1 and 2 was subjected to reverse phase HPLC. In the case of BEAK-tag-Aβ-P3*, the supernatant was incubated on ice for 2 h to allow fusion protein to aggregate. Precipitated protein was harvested by centrifugation (15000 RPM, 20 min, 4 °C), then resuspended in 5 % acetic acid and centrifuged again (as previously) in order to remove remaining impurities. Finally, aggregated fusion protein was dissolved in 0.1 M NaOH and subjected to trypsin cleavage.

**Reverse-phase HPLC**

BEAK-tag-TGFBIp1 and 2 fusion proteins before and after trypsin (or TEV) digestion were purified using reverse-phase HPLC system (Agilent) and a semi-preparative C8 column (Agilent). The proteins were eluted from the column with a linear gradient of acetonitrile containing 0.1 % TFA (trifluoroacetic acid) and freeze-dried to remove the solvents. The purity of the proteins was assessed by SDS-PAGE and their molecular weight verified by MALDI-TOF.

**Trypsin cleavage**

Purified BEAK-tag-TGFBIp1 and 2 fusion proteins were dissolved in the trypsin digestion buffer (50 mM Tris pH 8.9, containing 10 % acetonitrile) to obtain final protein concentration approx. 0.5 - 1.5 mg/mL. Trypsin powder (Trypsin Gold, MS grade, Promega) was dissolved in 50 mM acetic acid to the final concentration of 1 mg/mL and added to the reaction mixture in a 1:500 volume ratio (trypsin/protein). Enzymatic cleavage was carried out overnight at 37 °C. Reaction was stopped by lowering pH to 3.3 by addition of concentrated acetic acid. Purification of enzymatic cleavage products was carried out using reverse-phase HPLC as described above. Fractions containing cleaved peptides were freeze-dried and their molecular weight verified by MALDI-TOF.

Dissolved BEAK-tag-Aβ-P3* fusion protein in 0.1 M NaOH (for details see protein extraction) was diluted with a trypsin cleavage buffer to the final protein concentration of 1 mg/mL. Addition of basic protein solution caused slight rise of the buffer pH which was adjusted back to pH 8.9 with 1 M HCl. Trypsin cleavage was carried out as described above. Addition of acetic acid terminated the reaction and induced aggregation of cleaved Aβ-P3* peptide. The mixture was incubated for an extra 15 minutes at room temperature and centrifuged (15000 RPM, 15 min, 20 °C). Supernatant (containing cleaved soluble BEAK-tag) was discarded while pellet (containing aggregated Aβ-P3* peptide) was resuspended in 5 % acetic acid and centrifuged again (as previously). This step was repeated twice in order to remove traces of BEAK-tag and other components used for the enzymatic reaction. Finally, the Aβ-P3* peptide pellet was freeze-dried to remove remaining acetic acid. Molecular weight of obtained Aβ-P3* peptide was verified by MALDI-TOF.

**Matrix assisted laser desorption/ionization-time of flight (MALDI-TOF) mass spectroscopy**

MALDI-TOF was used to verify the molecular weight of purified fusion proteins and peptides obtained after the trypsin (or TEV) cleavage. Saturated solution of sinapic acid (dissolved in 50 % water, 50 % ACN and 0.1 % TFA) was used as a matrix. 1 µL of fusion protein/peptide (1 mg/mL) was mixed with the matrix, then applied onto a MALDI plate and left for 30 min to dry prior to the analysis. Measurements were performed on a MALDI-ToF Kratos Axima ToF2 instrument (Kratos-Shimadzu Biotech) equipped with the N2 laser (set at 337 nm bandwidth and 4 ns pulse width).

**Sodium dodecyl sulphate polyacrylamide gel electrophoresis (SDS-PAGE)**

Fusion protein samples were analysed by SDS-PAGE using Mini-PROTEAN TGX 4- 20 % Precast Gels (Bio-Rad). Precision Plus Protein Dual Color Standard (Bio Rad) was used as a reference. The gels were stained with Coomassie Brilliant Blue stain.

**NMR**

Lyophilized fusion proteins were dissolved in 10 mM acetic acid (pH 3.3) containing 5% $D_2O$ and 0.1 mM DSS prior the NMR experiments. $^{1}H$-$^{15}N$ HSQC spectra were recorded on a 600 MHz Bruker Advance III NMR spectrometer equipped with 5 mm z-gradient TXI cryoprobe operating at 298 K. Spectra were processed using TopSpin 4.0.6 software.

**TEM sample preparation and imaging**

Lyophilized TGFBIp1 and 2 peptides were dissolved in PBS buffer (pH 7.2) to a final concentration of 50 µM. Aβ-P3* was prepared by dilution of peptide stock solution (in 0.1 M NaOH) to final concentration of 50 µM in 20 mM HEPES, 100 mM NaCl, 2 mM TCEP (pH 7.5). The samples were incubated at 37 °C, with shaking (180 RPM) for 7 days (TGFBIp1 and 2) and for 60 hours (Aβ-P3*), respectively. Next, the samples were applied on copper-rhodium 400 mesh holey grids with 15 nm carbon coating (prepared in-house), incubated for 1 minute followed by negative staining with 2 % uranyl acetate for 1 minute and then air dried. The sample was then examined under FEI T12, 120 kV transmission electron microscope equipped with a 4K CCD camera (FEI) between 48000x to 68000x magnifications under low dose conditions.

**Thioflavin T (ThT) assay**

ThT assay was used to study the amyloid fibril formation of the recombinant TGFBIp2 peptide obtained from TEV cleavage, chemically synthesized TGFBIp2 peptide was used as a control. Thioflavin T (Sigma) was dissolved in 1X PBS buffer pH 7.4 to the final concentration of 20 µM. The peptides were dissolved in the same buffer to the final concentration of 50 µM. All experiments were carried out in triplicates using COStar 96 well assay plate with black flat bottom at 37 °C in BioTek CYTATION reader with orbital shaking before each measurement. The excitation and emission measurements were taken at 440 nm and 485 nm, respectively.

**SUPPLEMENTARY FIGURES**

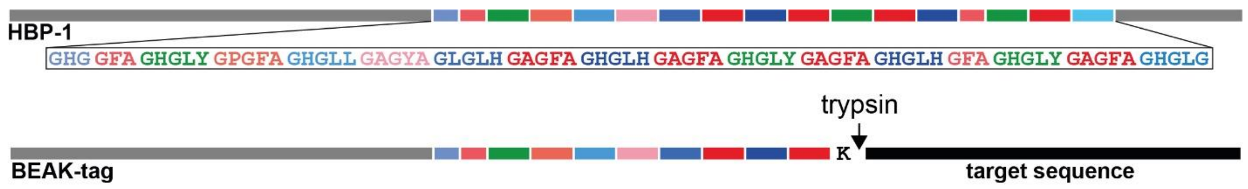


**Figure S1. BEAK-tag schematic illustration of the construct.** BEAK-tag was designed based on the HBP-1 sequence (upper panel, adapted from ref. 2). Modular region in the HBP-1 protein contains sequence responsible for initiating LLPS.[2] In the BEAK-tag (lower panel) part of the repetitive sequence was substituted with the target sequence, however the fusion protein retained its ability to undergo LLPS. Trypsin recognition site (K residue) was inserted between the BEAK-tag and the target sequence.

**a) eGFP**

MHHHHHHSSGVSKGEELFTGVVPILVELDGDVNGHKFSVSGEGEGDATYGKLTLKFICTTGKLPVPWPTLVTTLTYGVQ
CFSRYPDHMKQHDFFKSAMPEGYVQERTIFFKDDGNYKTRAEVKFEGDTLVNRIELKGIDFKEDGNILGHKLEYNYNSH
NVYIMADKQKNGIKVNFKIRHNIEDGSVQLADHYQQNTPIGDGPVLLPDNHYLSTQSALSKDPNEKRDHMVLLEFVTA
AGITLGMDELYKGGGS

**b) BEAK-tag**

MQLYGAPAVGGVVENAVNAAESGAAATHDAQGAYAEADTAGVLDVNHAEHHDGVHDASGYGFGGLAGHGGFAG
HGLYGPGFAGHGLLGAGYAGLGLHGAGFAGHGLHGAGFAK (ENLYFQG)*

**c) amyloidogenic peptides**

**TGFBIp1** DNQFS MLVAA IQSAG LTETL NR

**TGFBIp2** EPVAE PDIMD TNGVV HVITN VLQ

**Aβ-P3*** LVFFA EDVGS NHGAI IGLMV GGVV

**d) other peptides**

A$\beta_{1\text{-}15\ (5R/Q)}$ DAEFQ HDSGY EVHHQ

GY-23 GHGLY GAGFA GHGLH GFAGH GLY

Pep1 GHGLY GHGLY GHGLY GHGLY GHGLY GHGLY GHGLY GHGLY GHGLY GHGLY

Pep2 GHGLY GHGLY GHGLY GHGLY GAGFA GAGFA GHGLY GHGLY GHGLY GHGLY

Pep3 GHGLY GHGLY GAGFA GHGLY GAGFA GAGFA GHGLY GAGFA GHGLY GHGLY

Pep4 GHGLY GAGFA GAGFA GHGLY GAGFA GAGFA GHGLY GAGFA GAGFA GHGLY

A1H1$_{x3}$ GLGGYGGLYGGYPAATAVSHTTHHAPGGPAATAVSHTTHHAPGGPAATAVSHTTHHAPGLGGYGGLYGGY

**Figure S2. Protein and peptide sequences used in the study. a)** eGFP: enhanced green fluorescent protein, with N-terminal His-tag sequence (marked with blue) fused with a short linker (marked with green); a flexible GGGS linker (marked with green) was added to the C-terminus of the protein. **b)** BEAK: first 112 amino acids of HBP-1 protein (UniProtKB - A0A0G2UMW8, with methionine at the position 1 instead of the signal peptide), followed by lysine (K) residue (trypsin recognition site) or ENLYFQG (TEV recognition side), *used to prepare control construct BEAK-tag-TEV-TGFBIp2. **c)** Amyloidogenic regions from the transforming growth factor β-induced (TGFBI) protein: TGFBIp1 (512-533, 514R/Q) and TGFBIp2 (611-633, 620A/D). The peptides are highly important to understand the amyloid fibril formation with specific amino acid changes in that region of the protein. The 620A/D is a novel mutation in TGFBI gene (c.620 A>D) identified in a patient from Singapore.[3] Aβ-P3* engineered version of P3 (Aβ17-40, mutated residues marked in red). Mutation of basic residues were introduced to prevent cleavage with trypsin. **d)** other peptides expressed with the BEAK-tag: Aβ1-15 (5R/Q) fragment of amyloid β, GY-23 and Pep1-4 – repetitive sequences derived from the sequence of HBP-1 protein, A1H1$_{x3}$ amyloid-like β-sheets forming peptide (AATAVSHTTHHA) identified within the suckerin protein family flanked by flexible linker sequences.[4]

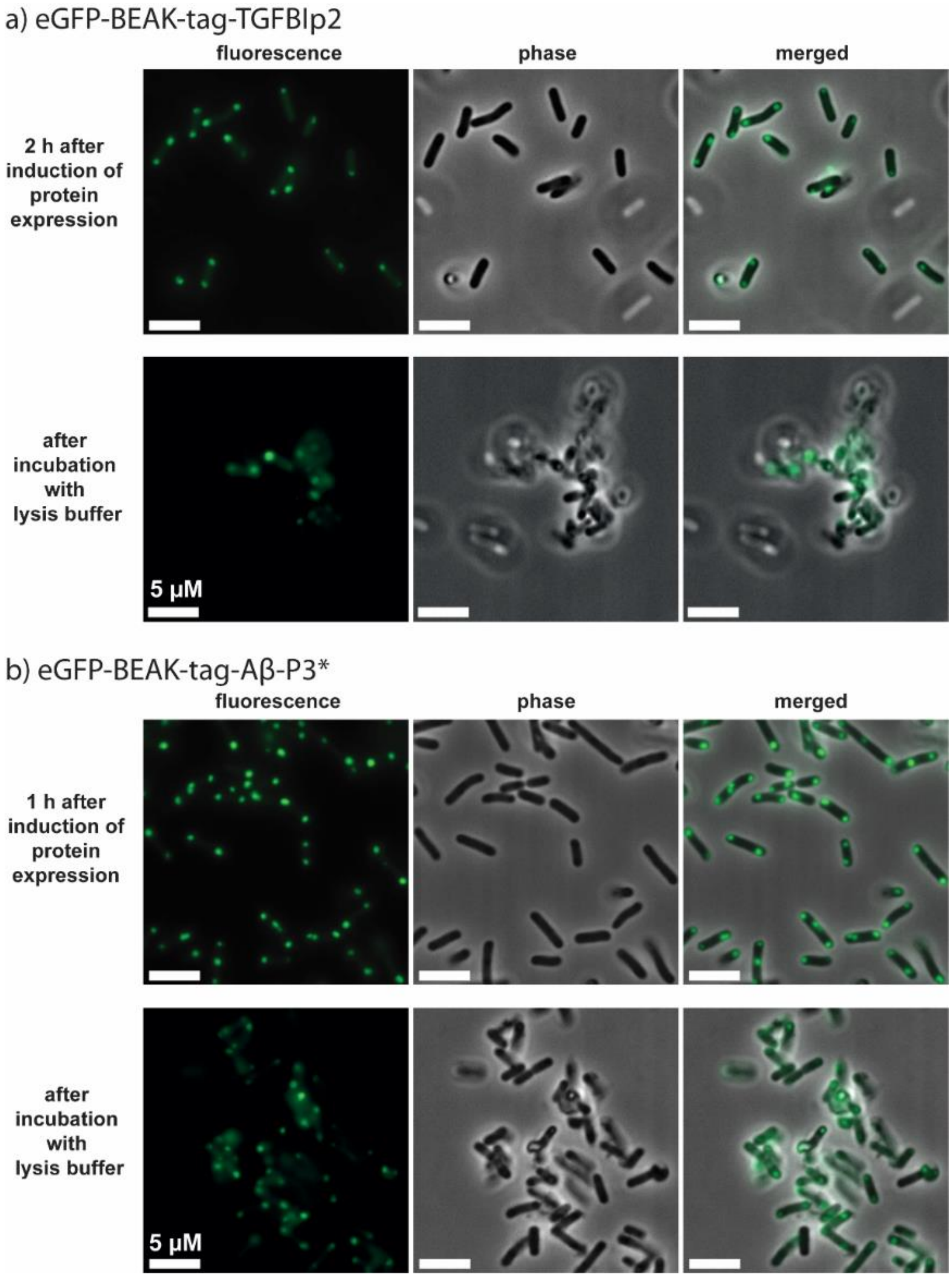


**Figure S3. Fluorescence microscopy and phase contrast images of bacterial cells expressing. a)** eGFP-BEAK-tag-TGFBIp2 and **b)** BEAK-tag-Aβ-P3* fusion proteins. Images recorded after induction of protein expression (upper panels) show fluorescent condensates localized near the pole regions of the cells. Incubation with lysis buffer (lower panels) leads to diffusion of the fluorescent signal indicating liquid-like properties of the protein condensates.

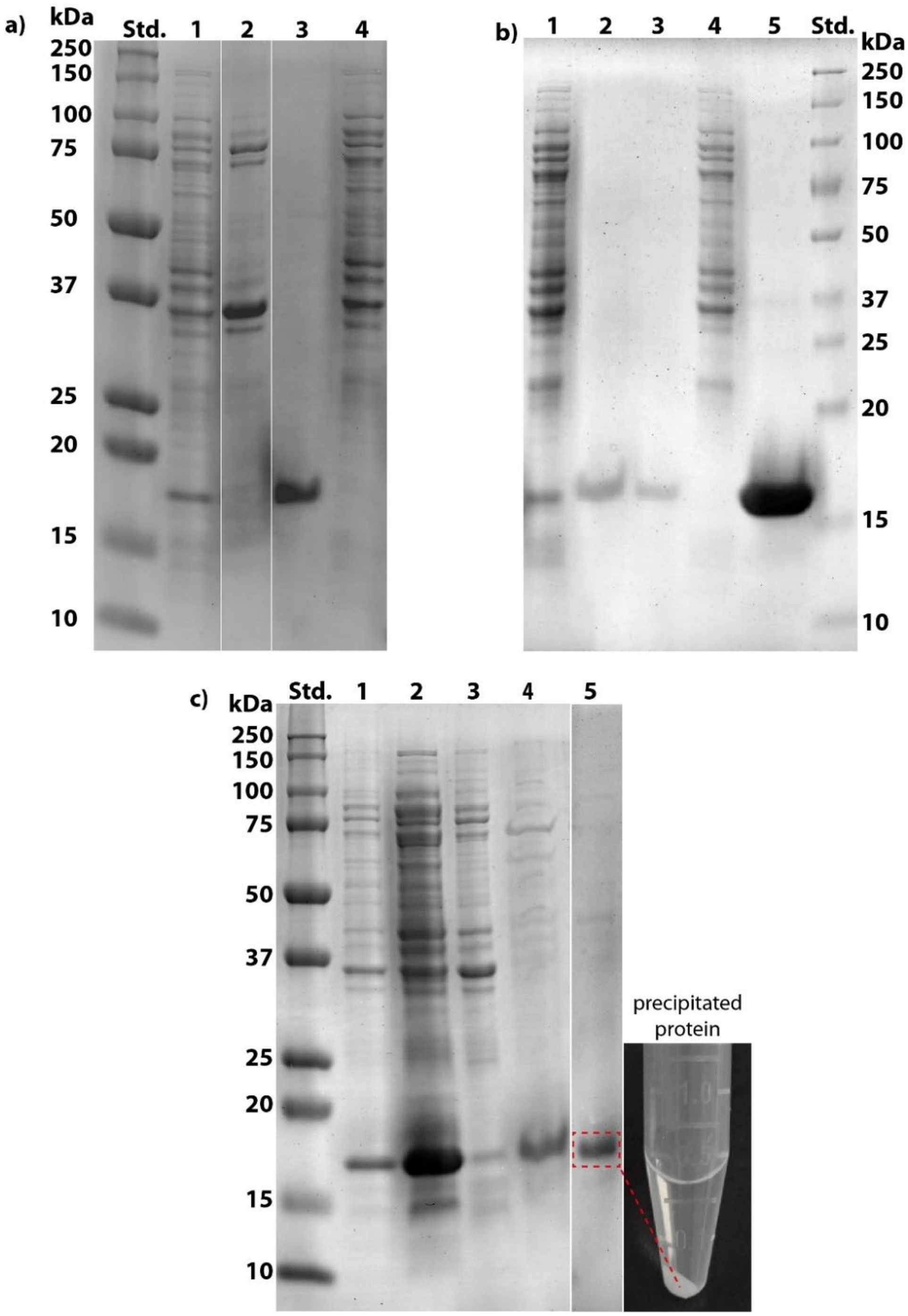


**Figure S4. BEAK-tag facilitates recombinant production of biologically functional aggregation-prone peptides.** SDS-PAGE **a)** BEAK-tag-TGFBIp1; Std.: protein standard; 1: soluble fraction after cell lysis; 2: insoluble fraction after cell lysis; 3: extracted proteins, 4: precipitated proteins during extraction **a)** BEAK-tag-TGFBIp2; 1: soluble fraction after cell lysis; 2, 3: extracted protein; 4: precipitated proteins during extraction; 5: purified protein after HPLC, Std.: molecular weight protein standard. **c)** BEAK-tag- Aβ-P3*; 1, 2: soluble fraction after cell lysis; 3: insoluble fraction after cell lysis, 4: extracted protein; 5: dissolved precipitated protein present in the picture.

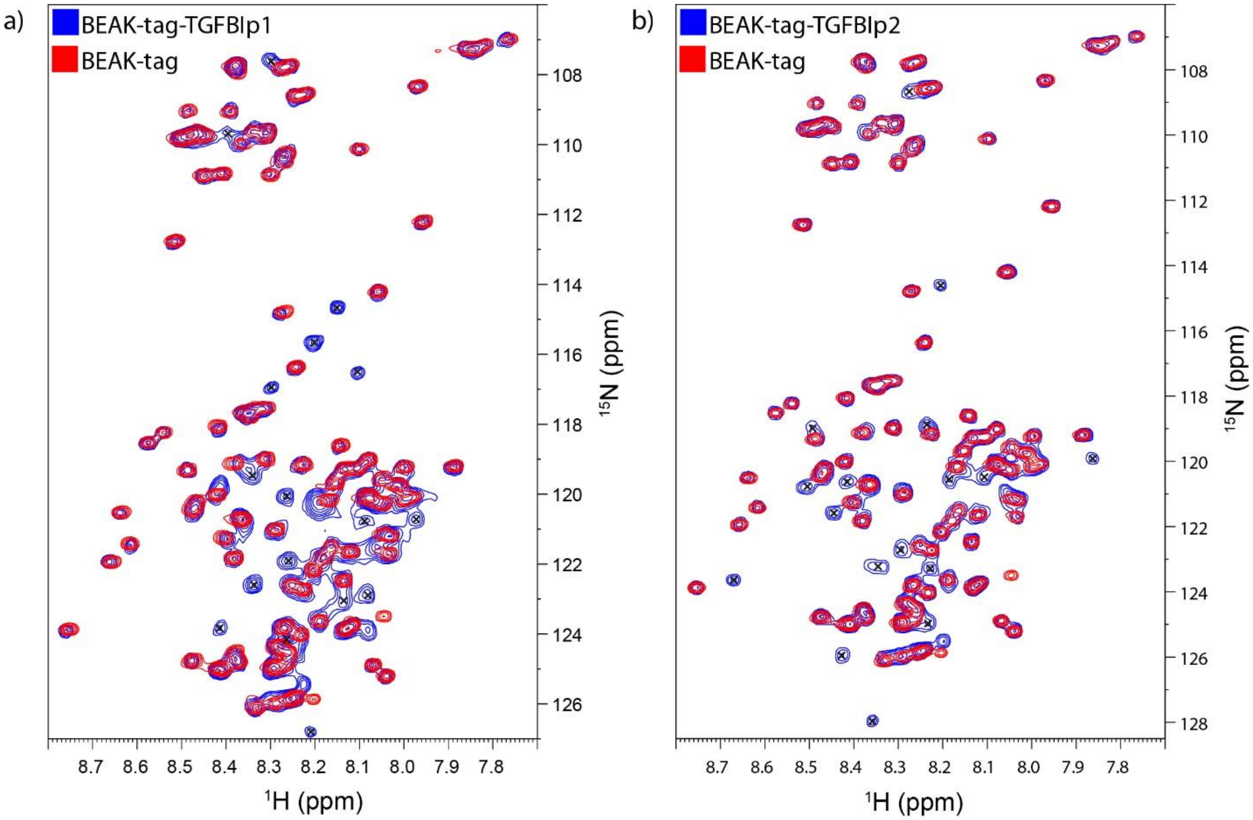


**Figure S5. Overlay of the 1H-15N-HSQC NMR spectra of the BEAK-tag (without fused peptide) a)** BEAK-tag-TGFBIp1 **b)** BEAK-tag-TGFBIp2 fusion proteins. Cross-peaks assigned to the TGFBIp1 and 2 peptides marked with "x". The chemical shifts of the target peptides in the fusion construct correspond to those observed in the synthetically produced version and exhibited uniform intensity, indicating the absence of secondary structure in the fused peptide as well as the random coil nature of the conformational ensemble.

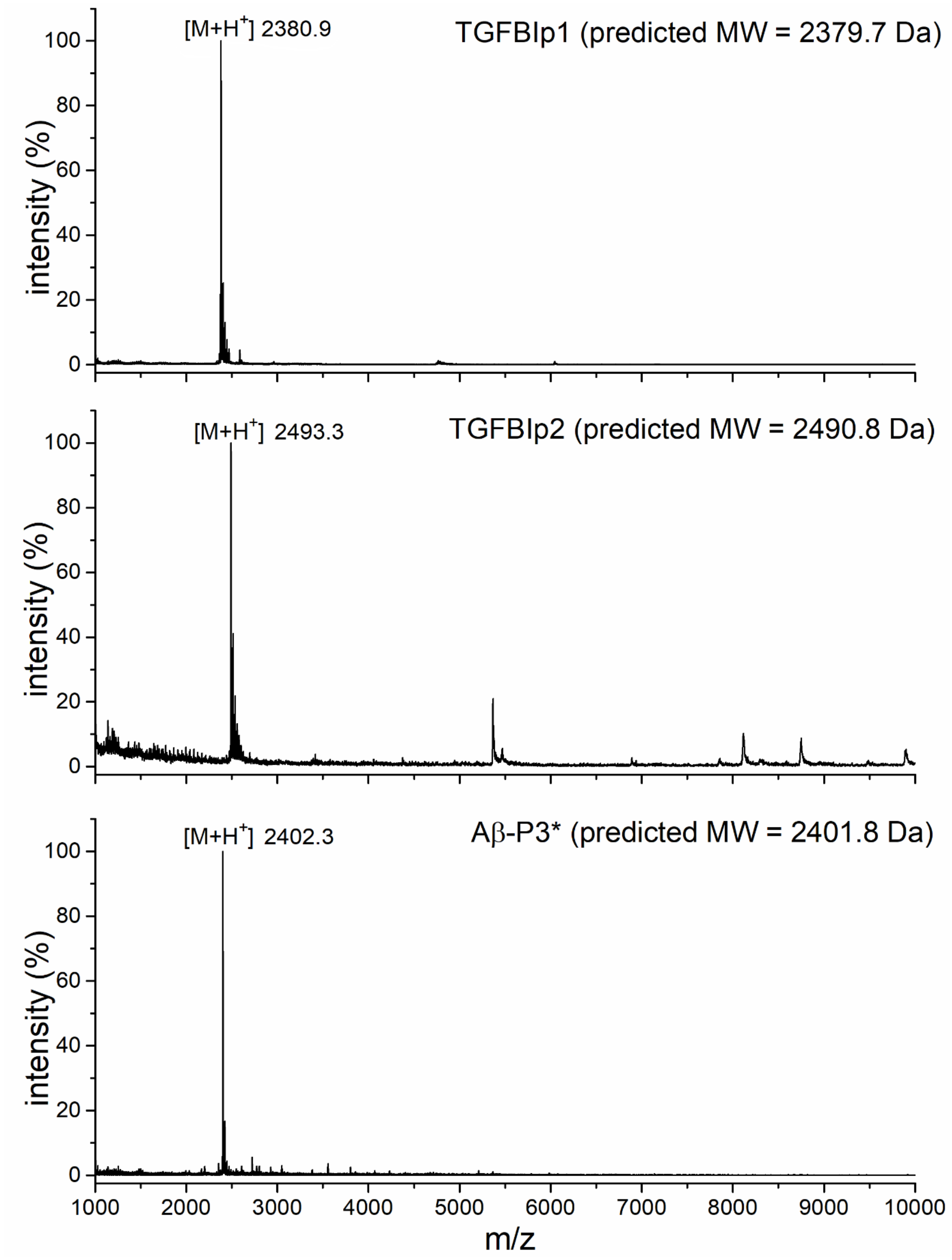


**Figure S6. MALDI-TOF spectra of purified amyloidogenic peptides.** Predicted molecular weight (MW) calculated using ProtParam tool available at https://web.expasy.org/cgi-bin/protparam/protparam.

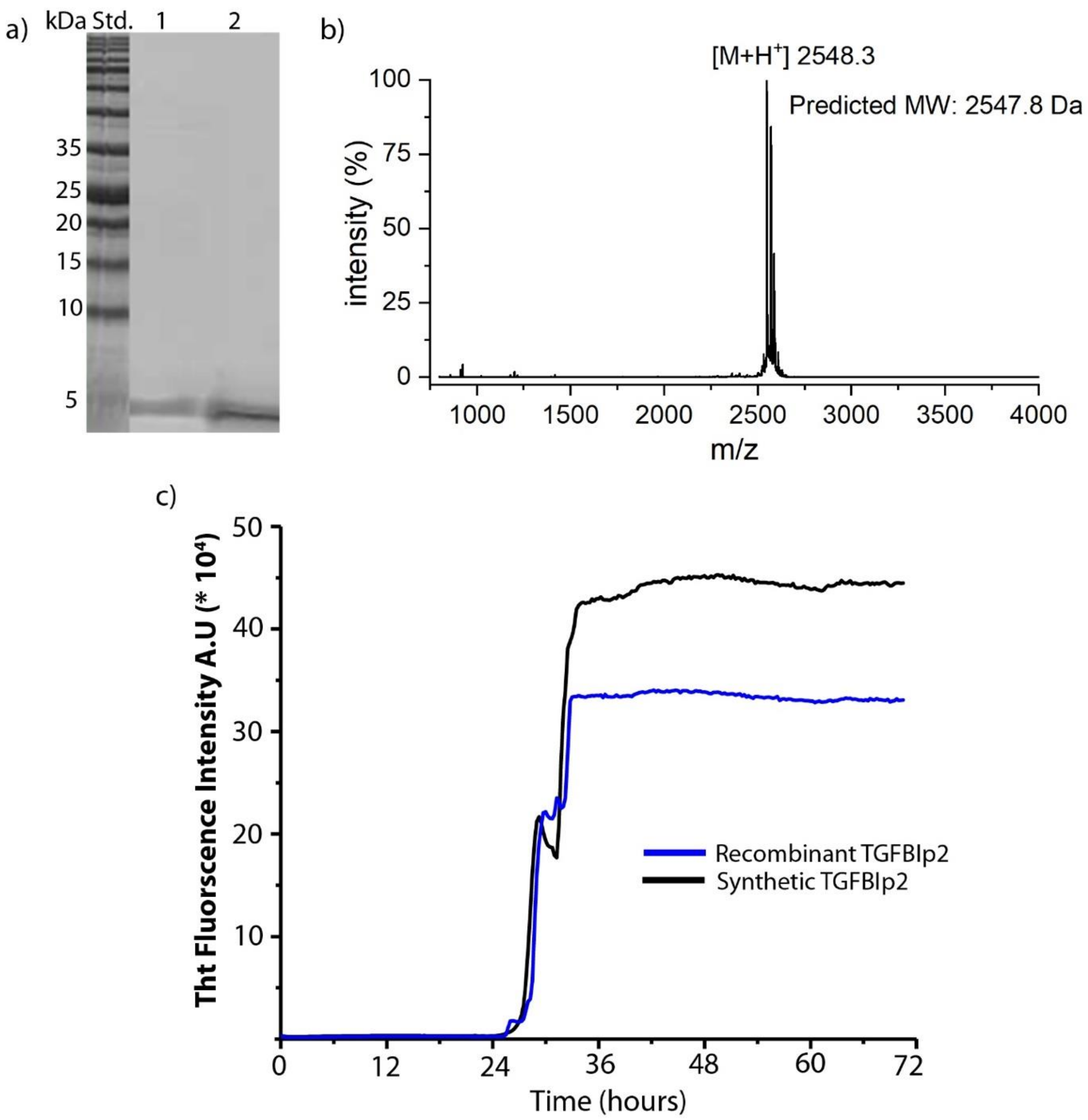


**Figure S7. Qualitative and functional analysis of the TGFBIp2 peptide obtain after TEV cleavage of the BEAK-tag-TEV-TGFBIp2 control protein. a)** SDS-PAGE showing purified TGFBIp2 peptide after reverse phase HPLC (line 1), synthetic TGFBIp2 used as a control (line 2), Std. – MW standard. **b)** MALDI-TOF spectrum of purified TGFBIp2 peptide (the peptide contains extra G residue remaining after TEV cleavage, thus its MW differ from the TGFBIp2 obtained from trypsin cleavage (Figure S6) that does not contain any extra residues). **c)** ThT assay showing amyloid formation of the recombinant TGFBIp2 peptide in comparison with its synthetic analogue. Both peptides are showing similar pattern of amyloid formation.

**REFERENCES**

[1] C. Engler, R. Kandzia, S. Marillonnet, PLoS One 2008, 3, e3647.

[2] B. Gabryelczyk, H. Cai, X. Shi, Y. Sun, P. J. M. Swinkels, S. Salentinig, K. Pervushin, A. Miserez, Nat. Commun. 2019, 10, 5465.

[3] R. Lakshminarayanan, E. N. Vithana, S-M. Chai, S. S. Chaurasia, P. Saraswathi, A. Venkatraman, C. Rojare, D. Venkataraman, D. Tan, T. Aung, R. W. Beuerman, J. S. Mehta British Journal of Ophthalmology 2011, 95, 1457–1462.

[4] S. H. Hiew, A. Sánchez-Ferrer, S. Amini, F. Zhou, J. Adamcik, P. Guerette, H. Su, R. Mezzenga, A. Miserez, Biomacromolecules 2017, 18, 4240–4248.
